# Accurate long-read transcript discovery and quantification at single-cell resolution with Isosceles

**DOI:** 10.1101/2023.11.30.566884

**Authors:** Michal Kabza, Alexander Ritter, Ashley Byrne, Kostianna Sereti, Daniel Le, William Stephenson, Timothy Sterne-Weiler

## Abstract

Accurate detection and quantification of mRNA isoforms from nanopore long-read sequencing remains challenged by technical noise, particularly in single cells. To address this, we introduce Isosceles, a computational toolkit that outperforms other methods in isoform detection sensitivity and quantification accuracy across single-cell, pseudo-bulk and bulk resolution levels, as demonstrated using synthetic and biologically-derived datasets. Isosceles improves the fidelity of single-cell transcriptome quantification at the isoform-level, and enables flexible downstream analysis. As a case study, we apply Isosceles, uncovering coordinated splicing within and between neuronal differentiation lineages. Isosceles is suitable to be applied in diverse biological systems, facilitating studies of cellular heterogeneity across biomedical research applications.

## Main Text

Alternative splicing (AS) contributes to the generation of multiple isoforms from nearly all human multi-exon genes, vastly expanding transcriptome and proteome complexity across healthy and disease tissues ^1^. However, current short-read RNA-seq technology is restricted in its ability to cover most exon-exon junctions in isoforms. Consequently, the detection and quantification of alternative isoforms is limited by expansive combinatorial possibilities inherent in short-read data ^2^. Short read lengths can impose additional challenges at the single-cell level. For example, nearly all isoform information is lost with UMI-compatible high-throughput droplet-based protocols which utilize short-read sequencing at the 3’ or 5’ ends ^3^. Recent advances in long-read sequencing technologies provide an opportunity to overcome these limitations and study full-length transcripts and complex splicing events at both bulk and single-cell levels, yet downstream analysis must overcome low read depth, high base-wise error, pervasive truncation rates, and frequent alignment artifacts ^4^. To approach this task, computational tools have been developed for error prone spliced alignment ^5^ and isoform detection/quantification ^6–13^. However, these tools vary widely in accuracy for detection and quantification, their applicability to bulk or single-cell resolutions, and in their capabilities for downstream analysis.

Here we present Isosceles (the ***Iso***forms from ***s***ingle-***ce***ll, ***l***ong-read ***e***xpression ***s***uite); a computational toolkit for reference-guided *de novo* detection, accurate quantification, and downstream analysis of full-length isoforms at either single-cell, pseudo-bulk, or bulk resolution levels. In order to achieve a flexible balance between identifying *de novo* transcripts and filtering misalignment-induced splicing artifacts, the method utilizes acyclic splice-graphs to represent gene structure ^14^. In the graph, nodes represent exons, edges denote introns, and paths through the graph correspond to whole transcripts (Fig 1a). The splice-graph and transcript set can be augmented from observed reads containing novel nodes and edges that surpass reproducibility thresholds through a *de novo* discovery mode, enhancing the adaptability of the analysis. In the process, sequencing reads are classified relative to the reference splice-graphs as either node-compatible (utilizing known splice-sites) or edge-compatible (utilizing known introns), and further categorized as truncated or full-length (Fig. 1a). Full-length reads can be directly assigned to known transcripts, meanwhile those representing novel transcript paths are assigned stable hash identifiers. These identifiers facilitate ease of matching *de novo* transcripts across data from the same genome build, irrespective of sequencing run, biological sample, or independent studies. In contrast, truncated reads may introduce ambiguity in terms of their transcript of origin, reflecting a challenge commonly found in short-read data analysis. To address this, we utilize a concept developed for short-read methods, Transcript Compatibility Counts (TCC) ^15^, as the intermediate quantification of all reads. TCCs are used to obtain the maximum likelihood estimate of transcript expression through the expectation-maximization (EM) algorithm (^16,17^; see Methods). This approach tackles another challenge: accurately quantifying transcripts at multiple single-cell resolution levels. First, transcripts can be quantified through EM within single-cells, which can be subsequently used to obtain a neighbor graph and low dimensional embedding (eg. with common tools like Seurat ^18^). Second, transcripts can be quantified at the pseudo-bulk level through EM on the TCCs summed within cell groupings (Fig. 1b). This configuration enables versatility of quantification; pseudo-bulk can be defined by the user in numerous ways, such as through marker labeling, clustering, windows along pseudotime, or for each cell based on its k-nearest neighbors (kNN). Downstream statistical analysis and visualization for percent-spliced-in and alternative start and end sites is seamlessly integrated to facilitate biological interpretation of isoforms.

**Figure 1.**
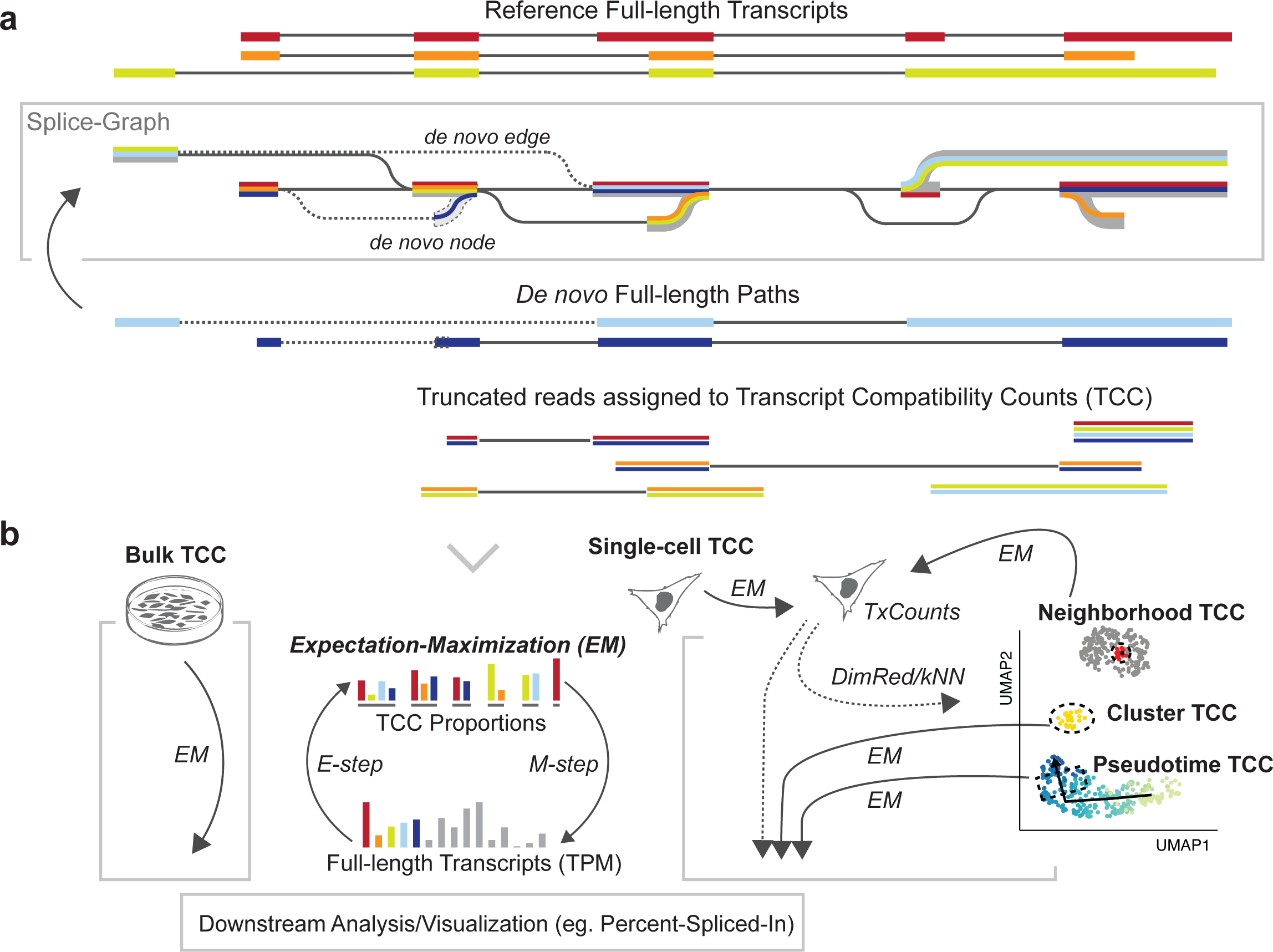
(a) Schematic of Isosceles splice-graph building and path representation of transcripts (colored lines). Augmentation with de novo nodes and edges (dashed). Ambiguous reads are assigned to TCCs to be quantified using the expectation-maximization (EM) algorithm (bottom > panel b). (b) The Isosceles approach to multi-resolution quantification using the EM algorithm. Transcripts quantified from single-cell TCCs using EM (grey cell, right) can be used for dimensionality reduction (DimRed) with UMAP or to derive a k-nearest neighbors graph (kNN). The original single-cell TCCs can be aggregated based on user-defined pseudo-bulk groupings and then transcripts re-quantified, either for clusters/markers, in windows along pseudotime or for each cell based on its neighborhood from kNN.

To robustly assess Isosceles performance against a wide-array of currently available software ^6–13^, we simulated ground-truth nanopore reads from reference transcripts proportional to the bulk expression profile of an ovarian cell line, IGROV-1, using NanoSim ^19^ (see Methods). In the evaluation of annotated transcript quantification against the ground-truth, Isosceles outperforms other programs, achieving a highly correlated Spearman coefficient of 0.96 (Fig. S1a). Bambu was the next best method at 0.92, while both IsoQuant and ESPRESSO were lower at 0.88.

Assessing quantification error through absolute relative difference, Isosceles decreases median and mean error by 21% compared to the next most accurate method, Bambu (0.23 vs. 0.29 and 0.41 vs. 0.52; Fig. 2a and S1a). Importantly, the reduction in error over other methods is even more pronounced, demonstrating ∼45% lower error than the median performer ESPRESSO, and 67-85% lower error than the worst performer NanoCount due to lack of detection of many simulated transcripts (Fig. 2a and S1a).

**Figure 2.**
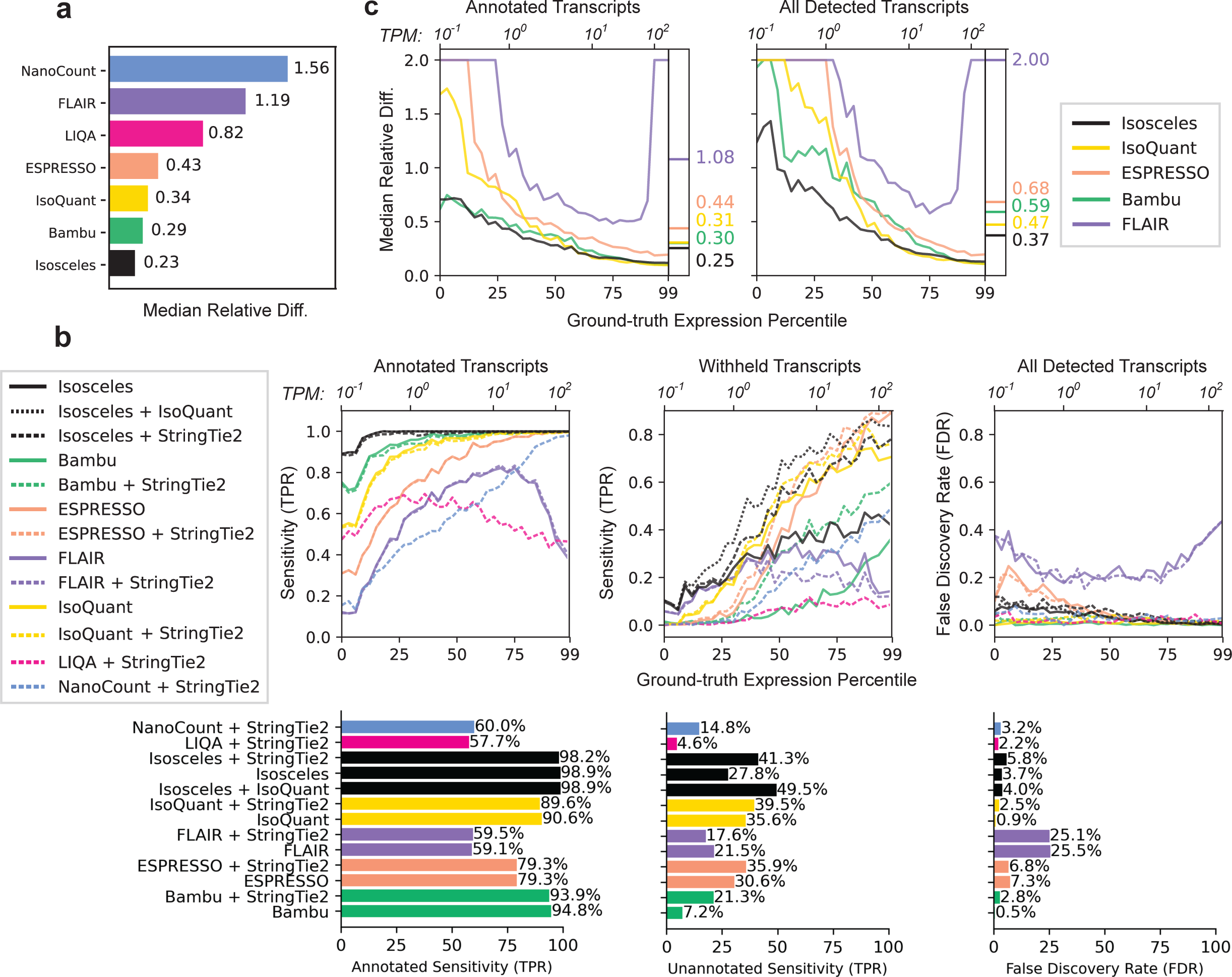
(a) Median relative difference of transcripts per million [TPM] values as defined by abs(ground_truth - predicted) / ((ground_truth + predicted)/2) for each method on reference transcripts. (b) Downsampling benchmarks for 30% transcripts withheld. Transcript detection defined as TPM > 0, the TPR and FDR detection rates as a function of the expression percentile (primary x-axis) and TPM values (secondary x-axis) of the simulated transcripts for single-program (solid) or pre-detection combinations (dashed), with overall TPR and FDR plotted as bars below the graphs. (c) Median relative difference of annotated and withheld transcripts (30% downsampling) as a function of the simulated expression level, as defined for panel b.

Since detection of both known and novel transcripts is a major attraction of long-read sequencing, we investigated the ability of various methods to detect 10%, 20% or 30% of transcripts when they are withheld from the annotation file (3269, 6537, 9801 transcripts respectively; 30% in Fig. 2b,c, 10% and 20% in Fig. S2a,b). Here, detection is defined as output of a transcript annotation with a splicing structure correctly matching a simulated transcript (irrespective of transcript start/end positions) and a quantification value greater than zero in transcripts per million (TPM > 0). We calculate the true-positive rate (TPR) as the number of correct transcripts detected from the total number with reads simulated and the false-discovery rate (FDR) as the percentage of incorrect transcripts out of the total detected. Notably, most methods output low TPR even for transcripts that are not withheld from the annotation file, so it is necessary to separate the TPR calculations for annotated and withheld transcripts (Fig 2b *left*, Isosceles=98.9% vs. median other=79.3%). Methods such as NanoCount and LIQA do not have a *de novo* detection mode, so we benchmark them with a pre-detection step using StringTie2 ^20^, adding this step to other tools for consistency (eg. Bambu, FLAIR, ESPRESSO, and also include IsoQuant alongside single-method detection for Isosceles; Fig. 2b *dashed lines*). While ESPRESSO and IsoQuant have modestly higher single-method TPR than Isosceles (2.8 and 7.8 percentage points respectively), the combination of Isosceles plus a pre-detection step with IsoQuant has the highest overall TPR across any program or combination thereof (Fig. 2b *middle*; 13.9 percentage points above IsoQuant alone). Importantly, Isosceles exhibits this relative gain in sensitivity at lower expression levels than other methods (<10 TPM). Overall, the resulting 49.5% TPR for Isosceles is obtainable at a reasonable FDR of 4.0%, which is comparable to other programs (Fig. 2b *right*; median FDR of 3.0%). When further considering the relative difference of quantification for annotated and withheld transcripts, Isosceles performs at 16.7% to 76.9% decrease in median error compared to other methods on annotated transcripts and 21.3% to 81.5% when including *de novo* (withheld) transcripts across the range of expression levels (Fig. 2c *left & right*; Fig. S3a). Similar to detection sensitivity, the most pronounced improvement in quantification accuracy occurs for the lowest half of expressed transcripts. Notably, while the single-program detection TPR of withheld transcripts in the latter comparison impacts on quantification accuracy, Isosceles alone still harbors less difference to ground-truth than other methods. These data suggest that state-of-the-art *de novo* detection and quantification can be achieved with Isosceles.

While known ground-truth values are effective for benchmarking performance, the analysis of true biological data introduces additional complexities that simulations may not fully capture. To address this, we benchmark each method’s fidelity of quantification for the same biological sample and ability to differentiate decoy samples across bulk and single-cell resolutions. We perform nanopore sequencing on 10X Genomics single-cell libraries from the pooling of three ovarian cancer cell lines, IGROV-1, SK-OV-3, and COV504, noting that the cells separate into three clusters by transcript expression and that each cluster corresponds to a separate genetic identity according to Souporcell ^21^ (Fig. 3a; see Methods). Conducting bulk nanopore sequencing in parallel on MinION and PromethION platforms, we investigate the consistency of those same cell lines as well as the ability to distinguish against four additional ovarian cancer cell lines sequenced as decoys, namely COV362, OVTOKO, OVKATE, and OVMANA. We find that Isosceles consistently maintains the lowest mean relative difference (24-43% less than other methods) and the highest Spearman correlation (0.87 for Isosceles vs. 0.75 for the next highest, Sicelore) amongst methods quantified on the same cell line in bulk and pseudo-bulk (Fig. 3b-c). We further find that this performance is recapitulated when comparing across technical runs, between platforms, and independent of the number of cells included or transcripts compared for IGROV-1 (Fig. S3b-c; Fig. S4b). To ensure the observed results reflect accuracy and not merely precision, we stringently consider the consistency of difference between matched and decoy comparisons. Here, Isosceles exhibits a 1.4- to 2.9-fold greater absolute difference using Spearman correlation and mean relative diff. respectively as compared to other methods (lower bound of 95% confidence interval, see Methods; Fig. 3b-c; Fig. S4a,c). To provide orthogonal support for this conclusion, we simulated a hundred cells at approximately ten thousand reads per cell using NanoSim with a single-cell error model (see Methods). While all methods show inflated error for single-cells compared to pseudo-bulk, Isosceles harbors lower average error than other methods for both, demonstrating quantification accuracy even in a data-sparse context (Fig. 3d).

**Figure 3.**
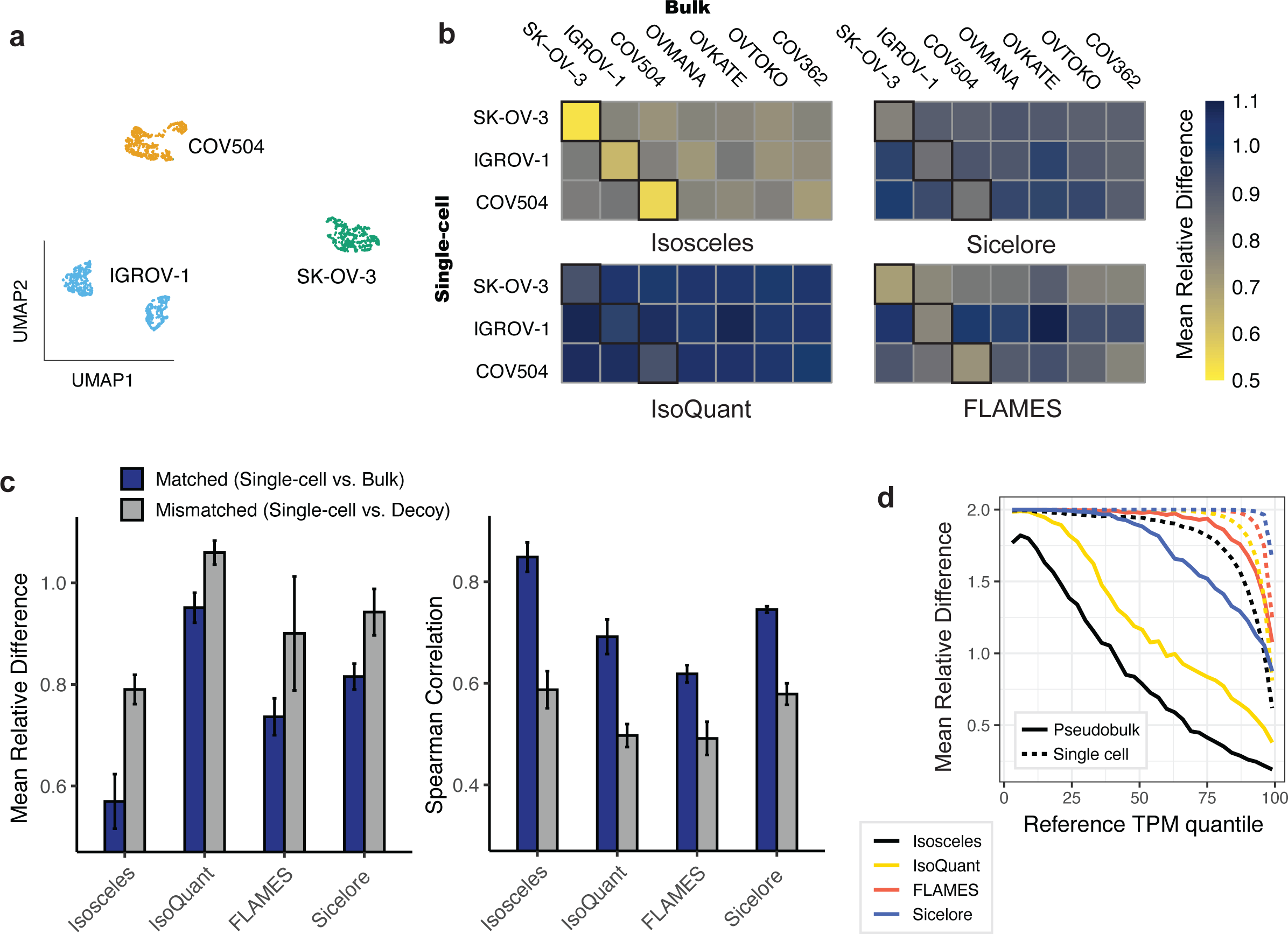
(a) 2D UMAP embedding of transcript-expression level quantifications from nanopore data of pooled IGROV-1, SK-OV-3, and COV504 ovarian cell lines, subsequently colored by genetic identity (according to Souporcell). (b) Mean relative difference (color scale) of each program’s quantifications across resolutions (pseudo-bulk vs. bulk data) for the top 4000 most variable transcripts. (c) Mean relative difference and Spearman correlation across matched and decoy comparisons for the top 4000 most variable transcripts (error bars show std. deviation) (d) Mean relative difference (as defined for Fig. 2a) between ground truth and estimated TPM values from simulated reads at pseudo-bulk (solid lines) and single-cell level (dashed lines).

Isosceles’ capabilities for accurate and flexible quantification also enhance downstream analysis and biological discovery. To demonstrate, we reanalyzed 951 single-cell nanopore transcriptomes from a mouse E18 brain. Investigating transcriptional markers (Fig. S5), we observe the major cell types identified in the original study using Sicelore^9^. Isosceles quantifications provide greater resolution however, separating differentiating glutamatergic neurons into two distinct trajectories instead of one (annotated here as T1 and T2), in addition to the single GABAergic trajectory using Slingshot^22^ (Fig. 4a). We also observe separation of radial glia and glutamatergic progenitor cells, which were connected in the original study. Isosceles’ versatility of pseudo-bulk quantification coupled to generalized linear models (GLM), further distinguishes downstream experimental design capabilities for biological discovery. For example, to investigate transcriptional dynamics within trajectories we apply the EM algorithm to pseudo-bulk windows, quantifying transcript expression as a function of pseudotime. To summarize individual transcript-features, Isosceles provides the inclusion levels of alternative splicing (AS) events, such as alternative exons and splice sites quantified as percent-spliced- in^2,23^ [PSI] or counts-spliced-in [CSI] (see Methods). In order to test for differential inclusion versus exclusion as a function of pseudotime (or any other condition), Isosceles seamlessly integrates with the DEXseq package to utilize GLMs in the context of splicing (see Methods).

**Figure 4.**
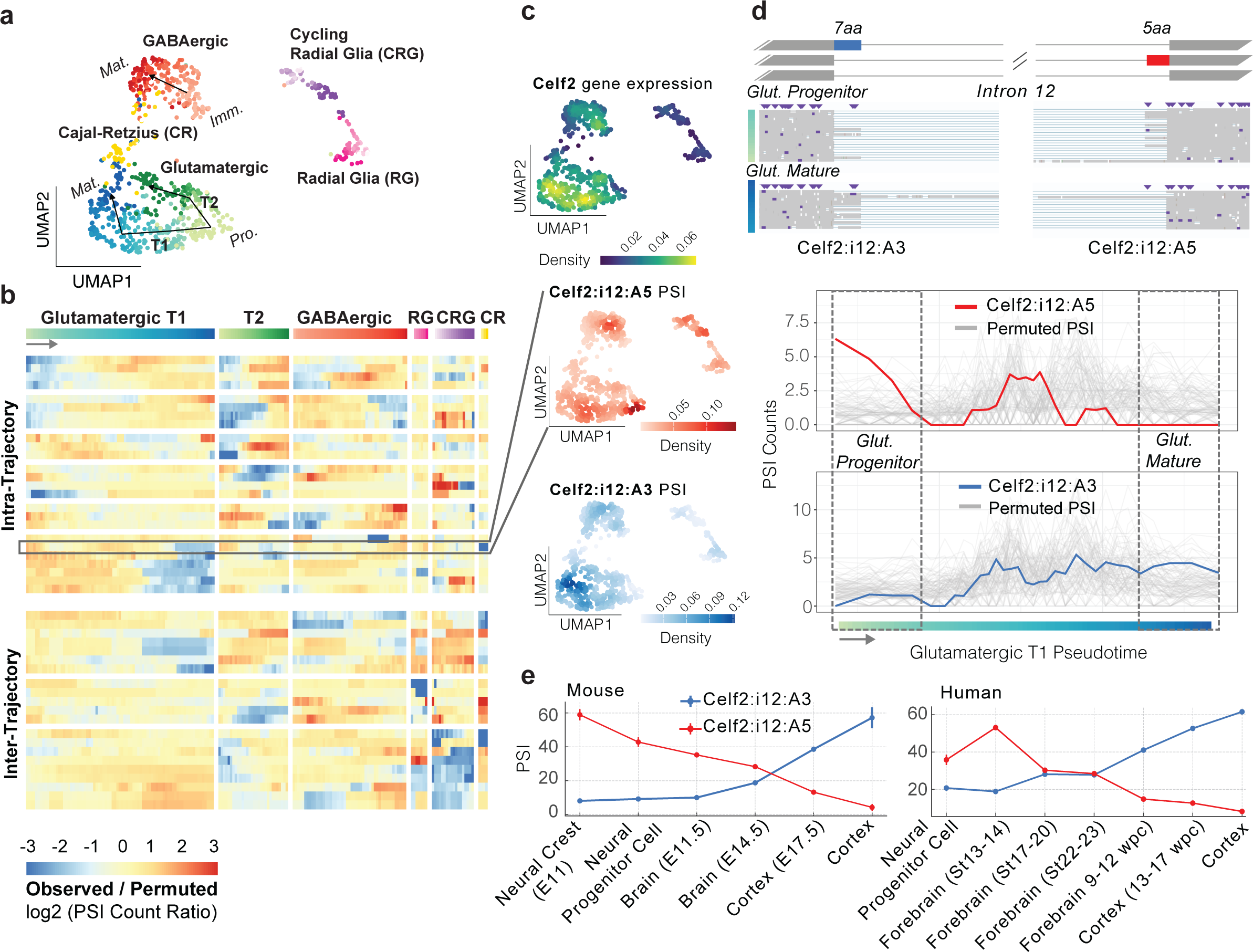
(a) 2D UMAP embedding from PCA performed jointly on variable gene and transcript features. Gradient coloring by pseudotime according to each trajectory. Glutamatergic progenitors are abbreviated Pro., Immature GABAergic neurons as Imm. and Mature neurons of both sub-types as Mat. T1 and T2 describe the two trajectories of Glutamatergic neurogenesis observed. (b) Heatmap of significant AS events colored by the ratio of observed CSI vs. permuted CSI. Permutations within (top) or across all (bottom) trajectories are separated. (c) UMAP density column from top to bottom: Celf2 gene expression, Celf2 alternative 5′ splice site (A5) in intron 12 (Celf2:i12:A5; chr2:6560659-6560670, row highlighted in panel b), and the juxtaposed alternative 3′ splice site (A3) for intron 12 (Celf2:i12:A3; chr2:6553965-6553982). (d) AS event diagram on the (top) of Celf2 gene intron 12 where exons are shown as boxes and introns as lines (gene on the ‘-’ strand), with the A5 event in red, the A3 event in blue, and reads from cells in the beginning and the end of the glutamatergic T1 trajectory shown respectively (from boxed regions annotated in the bottom panel). Bottom panel shows plots of CSI for windows along pseudotime for the observed data (A5, red) and (A3, blue) plotted over the background permutations in gray. (e) Mean PSI values with standard error as bars for human (left) and mouse (right) samples from the VastDB Mmu10 and Hsa38 short-read splicing databases^24^.

Applying the method identifies 25 AS events changing within trajectories as well as 21 changing between trajectories respectively (Table S1). Isosceles also implements the ‘isoform switching’ approach utilized in the original study (see Methods). However, we note that applying this method only identifies transcripts changing between major clusters, and none within glutamatergic or GABAergic neurogenesis trajectories (including the exemplar genes Clta and Myl6 presented in the original study; eg. Fig S6a).

One major challenge in the interpretation of single-cell data at the transcript-level (or event-level) is that fluctuations in detection or quantification may be attributable to gene expression changes alone. To decouple splicing dynamics and visualize them independently, we utilize a permutation-based approach. We estimate a background distribution by shuffling each gene’s splicing quantification among cells expressing that gene (within and between trajectories). We then visualize log ratios of the observed CSI values versus the mean expected CSI from these permutations (Fig. 4b; see Methods). Here, we observe AS events that exhibit precise changes within specific neuronal differentiation trajectories (such as only T1 or T2), including several RNA binding proteins (eg. Celf2, Hnrnpa2b1, Luc7l3, Ythdc1). Exemplifying a unique mode of alternative splicing in the gene Celf2, we observe a coordinated switch from one alternative donor splice site to an alternative acceptor splice site in the same intron as cells differentiate from glutamatergic progenitor to mature neurons (T1 trajectory, Fig. 4c-d). To validate the statistical significance of this event, we compare observed to permuted values using a stringent empirical test (see Methods). Here, we find the splicing-change is robustly independent of the overall changes in Celf2 expression that simultaneously occur (Fig. 4c-d & Fig. S7c; pval < 3.8x10^-4^). Underscoring biological significance, we note the two alternative splice sites have orthologs in other mammalian species (as annotated in VastDB ^24^) and high sequence conservation in the intronic region surrounding both splice sites (Fig. S7a-b). We validate the conserved mutual exclusivity and switch-like splicing change in human and mouse, recapitulating the longitudinal observation across embryonic brain samples from bulk short-read datasets ^24^ (Fig. 4e), including an *in vitro* study of mouse neuronal differentiation ^25^ (Fig. S7d).

In summary, Isosceles is a computational toolkit with favorable performance compared to other methods, as demonstrated through rigorous benchmarks on simulated and biological data from nanopore sequencing across ovarian cell lines. In these benchmarks, Isosceles performs transcript detection and quantification with accuracy, revealing improvements over existing methods that are most pronounced at lower expression levels. Notably, transcription factors and other regulatory proteins typically exhibit low gene expression levels, accompanied by rapid, fine-tuned regulation in mRNA and protein turnover rates ^26^. Such regulatory genes are frequently the focus of single-cell biological investigations, underscoring the importance of precision in this range. Through multi-resolution sequencing of ovarian cancer cell lines, we benchmark fidelity of quantification, demonstrating Isosceles’ performant capacity to consistently reproduce results for the same sample, and to differentiate among related yet distinct samples. Such intrinsic differences between cell lines, even those of the same tissue origin, may be more substantial than many biological changes typically investigated in biomedical research.

We further illustrate that these performant capabilities are enabling in the context of biological discovery. In our case study, we utilize Isosceles to uncover the dynamics of alternative splicing in differentiating neurons. Here, Isosceles provides enhanced resolution and reveals numerous AS events not reported in the original study. Importantly, these results reveal fine-tuned regulation within fate-determined trajectories and not only between major clusters (eg. radial glia vs. mature neurons). Among these events are genes encoding disease relevant RNA binding proteins that are themselves implicated in the regulation of neuronal differentiation. The Celf2 gene, for instance, plays a central role in neurogenesis, as it modulates the translation of target mRNAs through its shuttling activity ^27^. The example in Celf2 (presented in Fig. 4) highlights a switch-like splicing event that results in a conserved substitution of five to seven amino acids within the protein’s disordered region. This is akin to peptide changes introduced by microexons, which have been attributed functional roles in neurogenesis, including translational control of mRNAs through recruitment to membrane-less condensates, and dysregulation in disease ^28–30^. These results demonstrate that Isosceles is an effective method for hypothesis generation and biological discovery, offering insight into the splicing dynamics of a key regulator of differentiation in our case study.

Taken together, Isosceles is a flexible toolkit for the analysis of long-read bulk and single-cell sequencing that outperforms existing methods in detection and quantification across biological resolution levels. Based on its accuracy and flexibility for experimental designs, Isosceles will significantly aid researchers in transcriptomic studies across diverse biological systems.

## Data/Code Availability

Isosceles R package code, documentation, and vignettes are released on github (https://github.com/timbitz/Isosceles) under an open source GPL-3 license. All benchmarking code, virtual environments, and quantification data necessary to reproduce the figures/analyses in the manuscript are similarly released (analysis code: https://github.com/timbitz/Isosceles_paper, singularity containers: https://doi.org/10.5281/zenodo.8180648, benchmark quantifications: https://doi.org/10.5281/zenodo.8180604, raw simulated data: https://doi.org/10.5281/zenodo.8180695, mouse E18 brain scRNA-Seq data: https://doi.org/10.5281/zenodo.10028908). All biological sequencing data is deposited in the NCBI Gene Expression Omnibus (GEO) under GSE248118.

## Author Contributions

TSW and MK conceived of and designed the software methodology and computational experiments with contributions from the other authors. MK implemented the Isosceles package and both MK and AR performed benchmarking analyses. MK and TSW designed and performed the case study. KS performed the cell culture and AB and WS performed the sequencing protocols with preliminary analyses from DL. TSW wrote the manuscript with contributions from MK and all other authors.

## Acknowledgements

We would like to thank Bo Li, Hector Corrada-Bravo, William Forrest, John Marioni, Luca Gerosa, Marc Hafner, and Robert Piskol for helpful suggestions and feedback.

## Online Methods

### Isosceles Splice-graphs

Splice-graph compatibility is defined for reads using various stringency levels to match their concordance with existing knowledge. Reads are classified based on compatibility as Annotated Paths (AP), Path Compatible (PC), Edge Compatible (EC), Node Compatible (NC), De-novo Node (DN), Artifact Fusion (AF), Artifact Splice (AS), and Artifact Other (AX). AP refers to full-length transcript paths that perfectly match a reference transcript from the input gene annotation and are quantified by default. PC reads follow transcript paths that are a traversal of an AP, and may be truncated or full-length or with differing transcript start or end positions. EC reads traverse annotated splice-graph edges (introns) and may be truncated or full-length. NC reads are paths that traverse only annotated splice-graph nodes (splice-sites) but contain at least one novel edge. DN reads have paths that traverse a *de novo* node (splice-site). AF reads traverse paths connecting at least two splice-graphs for annotated genes that do not share introns with each other. AS reads are assigned to genes, but traverse an unknown and irreproducible node (splice-site), while AX reads lack compatibility due to ambiguous strand or lack of gene assignment.

Reads are also classified based on their truncation status, which includes Full-Length (FL), 5’ Truncation (5T), 3’ Truncation (3T), Full-Truncation (FT), and Not Applicable (NA). AP transcripts are automatically annotated as FL, and truncation status is checked only for PC, EC, NC, and DN transcripts. AF, AS, and AX transcripts are automatically labeled NA. Reference transcripts used for truncation status classification are recommended to be filtered to only the GENCODE ’basic’ dataset (tag=‘basic’), but also could be all transcripts in the provided annotations, as decided by the user. Full-length reads are those whose paths splice from a first exon (sharing a reference transcripts first 5’ splice site) and whose paths splice to a last exon (sharing a reference transcripts final 3’ splice site).

To add nodes with one or more de novo splice sites to the splice-graph, each splice-site must meet two conditions: it is observed in at least the minimum number of reads (default: 2) and it is connected to a known splice site in the splice-graph with least a minimum fraction (default: 0.1) of that known splice site’s connectivity. Additionally, annotations for known transcripts and genes are merged and extended based on specific criteria. For example, any annotated genes sharing introns with each other are merged into one gene and given a new gene_id & gene_symbol (comma-separated list of original Ensembl IDs and gene symbols). Annotated spliced (and unspliced) transcripts sharing the same intron structure, as well as transcript start and end bins (default bin size: 50 bp) are merged together and given a unique transcript identifier.

The method offers three modes of extending annotations to include *de novo* transcripts: *strict*, *de_novo_strict*, and *de_novo_loose*. In the *strict* mode, only AP transcripts are detected/quantified. In the *de_novo_strict* mode, AP transcripts and filtered FL transcripts of the EC and NC classes are included in quantification. In the *de_novo_loose* mode, AP transcripts and filtered FL transcripts of the EC, NC, and DN classes can be included.

For downstream analysis of individual transcript features, AS events are defined as the set of non-overlapping exonic intervals that differ between transcripts of the same gene. These are quantified as percent-spliced-in or counts-spliced-in according to the sum of the relative expression or the raw counts of the transcripts that include the exonic interval respectively. AS events are classified into different types similar to previous methods analyzing splicing from short-read data ^2^, including core exon intervals (CE), alternative donor splice sites (A5), alternative acceptor splice sites (A3), and retained introns (RI). Isosceles can also quantify tandem untranslated regions in the first or last exons including transcription start sites (TSS) and alternative polyadenylation sites (TES).

### Isosceles Quantification

We use the Expectation-Maximization (EM) algorithm to obtain the maximum likelihood estimate (MLE) of transcript abundances, as used previously in transcript quantification methods for short-read data such as our prior software Whippet ^2^, or the approach’s conceptual precursors RSEM ^16^ and/or Kallisto ^17^. Specifically, we quantify transcript compatibility counts (TCCs) based on fully contained overlap of reads to the spliced transcript genomic intervals (including an extension [default: 100 bp] for transcript starts/ends), with strand for unspliced reads ignored by default. For computational efficiency, TCCs matching more than one gene are disallowed in the current version. The likelihood function models the probability of observing the data given the current estimates of compatible transcript abundances, and is defined as described previously for transcript estimation from short-read data with Whippet ^2^, with the exception of effective transcript length. Here, due to the long length of nanopore reads, we define the effective transcript length to be the maximum of the mean read length vs. the transcript’s actual length, then divided by the mean read length. This directly accommodates shorter transcripts which would be fully spanned by the average read and are thus assigned an effective length of 1.0, whereas longer transcripts are represented proportionally to that value. In contrast, the user defined parameter specifying single-cell data does not use length normalization due to the anchoring of reads to the 5’ or 3’ ends of transcripts which assumes read coverage irrespective of transcript length. The EM algorithm iteratively optimizes the accuracy of transcript abundance estimates derived from TCCs, continuing until the absolute difference between transcript fractions is less than a given threshold (default:0.01) between iterations, or until the maximum number of iterations is reached (default: 250).

### Simulating ONT data

In this study, the Ensembl 90 genome annotation (only transcripts with the GENCODE ’basic’ tag) was used for all simulations, focusing specifically on spliced transcripts of protein-coding genes to exclude single-isoform non-coding genes. In order to simulate data with realistic transcriptional profiles, we quantified the expression of reference annotations in IGROV-1 cells using publicly available short-read data ([sample, project] accession ids: [SRR8615844, PRJNA523380]; https://www.ebi.ac.uk/ena/browser/view/SRR8615844) and Whippet v1.7.3 using default settings. Only transcripts with non-zero expression in IGROV-1 were retained for simulations. For detection benchmarks, the Ensembl 90 annotation file (in Gene Transfer Format [GTF]) was randomly downsampled such that the longest transcript of each gene was always retained to ensure at least one full-length major isoform for each gene (by 10%, 20%, and 30% downsampling, where 99.8-100.0% of downsampled transcripts had unique exon-intron architectures). In order to simulate Oxford Nanopore Technologies (ONT) reads using NanoSim, we trained error models on bulk nanopore RNA-Seq FASTQ files concatenated from sequencing three cell lines: SK-OV-3 (SAM24385455), COV504 (SAM24385457), and IGROV-1 (SAM24385458). Nanopore single-cell RNA-Seq (nanopore scRNA-Seq) read models were also generated from the pooled set of the aforementioned cell lines (SAM24404003). A total of 100 million reads were simulated from each error model and then the first 12 million reads deemed alignable by NanoSim were extracted.

To align the simulated reads provided in BAM format to all benchmark programs, Minimap2 was employed, using Ensembl 90 introns given in a BED file and applying a junction bonus parameter of 15. For the scRNA-Seq ONT dataset used to create the read model, various tools detected a similar number of cells (∼2460), but the median number of unique molecular identifiers (UMIs) per cell differed. The Sicelore preprocessing of ONT scRNA-seq, identified between 3,000 and 6,000 UMIs per cell, which were provided in BAM format for biologically derived data benchmarks to Sicelore, IsoQuant, and Isosceles with cell barcode and UMI tags annotated (Fig. 3a/b). In contrast, FLAMES, with its own UMI detection and deduplication processes, detected around 13,500 UMIs per cell. To strike a balance between the varying results from different tools, a compromise of 10,000 reads per cell was chosen for this study.

To simulate scRNA-Seq ONT data, a BAM file containing aligned simulated reads from the scRNA-Seq read model was randomly downsampled 100 times using samtools, with a subsampling proportion of 0.000833. This resulted in approximately 10,000 reads out of the original 12 million for each BAM file. A custom Python script (see supplemental Benchmark commands) was used to assign unique cell barcode sequences and UMI sequences for each read within the 100 BAM files. These subsampled BAM files were then merged and sorted using samtools.

### Biological data processing

The bulk RNA-Seq data included Promethion data (NGS3273), featuring eight sequencing libraries for seven ovarian cancer cell lines (OVMANA, OVKATE, OVTOKO, SK-OV-3, COV362, COV504, and IGROV-1), as well as two technical replicates for IGROV-1. For MinION platform data (NGS3082), two technical replicates for IGROV-1 were sequenced. Factors such as RAM performance and program speed determined the number of reads simulated in bulk simulations and downsampled in bulk data. For example, for performing cross platform correlations, the Promethion data was downsampled to 5 million reads to make it more comparable to MinION (∼6-7 million raw reads) and pseudo-bulk scRNA-Seq (3.5-4.5 million UMIs per cluster, as detected by Isosceles) in terms of total read depth. This decision was also influenced by an issue with IsoQuant (https://github.com/ablab/IsoQuant/issues/69), which limited its ability to process large read files in our hands. Notably, this issue persisted on a cluster node with 20 CPUs of 2.4GHz and allocated 230 GB of RAM.

The scRNA-Seq data (SAM24404003) consisted of a mix of three cell lines (SK-OV-3, COV504, and IGROV-1). The Illumina sequencing (LIB5445371_SAM24404003) was preprocessed using CellRanger (Version 6.0.1). The ONT sequencing (LIB5445493_SAM24404003) was preprocessed using the Sicelore workflow, resulting in a BAM file with cell barcode and unique molecular identifiers annotated.

All reads were aligned to the reference genome using minimap2 as discussed for simulated data. Mitochondrial transcripts common to all method’s output were removed, as they were strong outliers across methods. Additionally, three specific transcripts outliers across methods were removed: ENST00000445125 (18S ribosomal pseudogene), ENST00000536684 (MT- RNR2 like 8), and ENST00000600213 (MT-RNR2 like 12).

### Analysis of biological data

The correlation and relative difference analyses (Fig. S3b) compared annotated transcripts between bulk RNA-Seq data from two Promethion and two MinION sequencing replicates of IGROV-1, both within each platform (using replicates) and between platforms (using averaged data for each platform). For each comparison, only transcripts with a mean expression of at least 1 TPM were used. In Fig. S3c, scRNA-Seq and bulk RNA-Seq data were also compared, again considering only annotated transcripts. For each program, the IGROV-1 scRNA-Seq pseudo-bulk cluster (according to genetic identity from Souporcell) was compared with the averaged bulk RNA-Seq IGROV-1 expression values from two replicates for each platform.

Analyses were also restricted to transcripts with an expression of at least 1 TPM in the single-cell RNA-Seq results. Comparisons were made for each platform using top k cells (highest UMI count) using the top 5000 transcripts (highest mean expression) to ensure a comparable number of transcripts across software package, and top N transcripts (highest mean expression) for 64 top cells (highest UMI count) (Fig. S3c).

For Fig. 3a, scRNA-Seq and bulk RNA-Seq data analysis was conducted using Bioconductor packages (scran, scater, etc.) on the transcript and gene level for cells with at least 500 genes, considering 4000 top highly variable genes/transcripts. Heatmaps were generated to show correlations and mean relative difference between scRNA-Seq pseudo-bulk results for three cell line clusters and Promethion bulk RNA-Seq results for 7 ovarian cancer cell lines, similarly only including annotated transcripts. IGROV-1 expression was averaged from two replicates. To compare difference between matched and decoy metrics (Spearman correlation and mean relative difference), we calculate the absolute difference and compute the lower bound of the 95% confidence interval from the propagated error (as |x-y| - sqrt(sd(x)^2 + sd(y)^2) * 1.96).

For the case-study in Fig. 4, the raw reads were pre-processed to identify cell barcodes (CB) and unique molecular identifiers (UMI) according to the Sicelore workflow. The reads were subsequently aligned to the reference genome mm10/GRCm38 (with annotations derived from GENCODE M25), using Minimap2 with a junction bonus of 15, which targeted both annotated introns from Gencode M25 and those extracted from the VastDB mm10 GTF file ^24^. The aligned reads with CB and UMI annotations were subsequently quantified with Isosceles. The 951-cell dataset was filtered to exclude cells that expressed fewer than 100 genes. For dimensionality reduction, we combine Isosceles gene and transcript counts, culminating in the total identification of 3,760 variable features (with a target of 4,000), comprising 1,735 genes and 2,025 transcripts. We applied Principal Component Analysis (PCA), calculating 30 components using the scaled expression of the variable features. Cells were clustered using Louvain clustering (with resolution parameter of 2) on the Shared Nearest Neighbor (SNN) graph (setting a k-value of 10). The clusters’ identities were determined through gene set scores, particularly the mean TPM values of markers delineated in the original study (see Fig. S5). Additional marker genes were identified via the scran::findMarkers function requiring the t-test FDR to be significant (q-value < 0.05) in at least half of the comparisons to other clusters (selecting top 5 markers of each cluster).

Pseudotime analysis was performed using Slingshot for differentiating glutamatergic neurons (identifying two trajectories, T1 and T2), differentiating GABAergic neurons, radial glia, cycling radial glia and Cajal-Retzius cells (with one trajectory each). To implement the original ‘isoform switching’ analysis, pairs of clusters were compared, detecting marker transcripts through the specific scran::findMarkers function (Wilcoxon test). We filter for transcripts of the same gene showing statistically significant differences in opposite directions (i.e. one upregulated in one cluster, the other in another cluster). To analyze splicing changes within each trajectory, we used Isosceles to calculate aggregated TCC values for windows along pseudotime, defining the window size as 30 cells and the step size as 15 cells. AS events from variable transcripts abiding by further criteria were selected for downstream analysis. First, mean PSI values across all cells from the trajectory were between 0.025 and lower than 0.975 to exclude constitutively included/excluded events. Second, at least 30 cells must have values not equal to 0, 1, or 0.5, and 30 cells must have a value above 0.1 to select against events with only low counts.

Redundant PSI events, identical in read counts profiles within a trajectory, were excluded, and those with >0.99 spearman correlation were excluded from visualization in Fig. 4b and Fig. S6b. For comparative analysis, percent-spliced-in (PSI) count values are denoted as counts-spliced-in (CSI) and defined by PSI * gene counts. These are juxtaposed with exclusion PSI counts, calculated as [ (1 - PSI value) * gene counts ] and the inclusion/exclusion pair input into DEXSeq^31^. For each intra-trajectory comparison, our experimental design encompassed ‘∼sample + exon + pseudotime:exon’. Meanwhile, the inter-trajectory analysis included all trajectories with a design of ‘∼sample + exon + pseudotime:exon + trajectory:exon’, compared against a null model of ‘∼sample + exon’ using the LRT test.

To determine ratios of observed vs. expected CSI, we shuffle TCCs across cells with non-zero counts and apply the EM algorithm, calculating PSI for each window. To obtain expected CSI we multiply the shuffled PSI values * observed gene counts. The permutations are conducted for each AS event across 100 bootstraps. For empirical statistical validation of changes between the first and last windows of a trajectory (eg. for Celf2), we fit a negative binomial distribution to each window using maximum likelihood estimation (‘fitdistrplus’ package) on the permuted CSI, and calculate high and low one-tailed p-values for the observed CSI. Combining the high and low, and low and high p-values of the first and last windows respectively using fisher’s method, we defined an overall p-value as two times the minimum combined p-value. Specifically for heatmap visualization, a broad window size of 100 cells for glutamatergic & GABAergic neurons, and 50 cells for glia and CR cells, with a consistent step size of 3 cells for smoothing was utilized. The heatmap values were given as the log2 ratio of observed to expected, with a pseudocount of 0.1, defining the ratio between PSI counts and the average of the corresponding permuted PSI counts.

Benchmark command summary: https://github.com/timbitz/Isosceles_Paper/blob/devel/Benchmark_commands.md

### Software versions

| Software | Version |
| --- | --- |
| Isosceles | v0.0.3 |
| flair | v1.7.0 |
| stringtie | v2.2.1 |
| isoquant | v3.0.3 |
| NanoCount | v1.0.0.post6 |
| Sicelore | v2.0 |
| bambu | v3.2.5 (R 4.3.0, Bioconductor 3.17) |
| FLAMES | v0.1 |
| ESPRESSO | beta1.3.0 |
| nanosim | v3.1.0 |
| minimap2 | v2.24-r1122 |

### Cell culture

All cell lines used in this study were validated by STR analysis and verified mycoplasma negative by PCR. IGROV1, SK-OV-3, OVTOKO, OVKATE and OVMANA cell lines were cultured in RPMI-1640 supplemented with 10% heat-inactivated fetal bovine serum (FBS) and 2mM L-Glutamine. COV362 and COV504 cells were cultured in DMEM supplemented with 10% FBS and 2mM L-Glutamine. Cells were cultured in 37°C and 5% CO_2_ in a humidified incubator. Cell line source and catalogue numbers are provided in the table below. Cells were cultured in 10cm^2^ plates until they reached ∼60-80% confluency. For bulk analysis, RNA was purified using Qiagen’s RNeasy Plus Mini kit (Cat. #74134) according to manufacturer’s instructions. For single-cell analysis, IGROV1, SK-OV-3 and COV504 cells were trypsinized and pooled together at a 1:1:1 ratio at a concentration of 1000 cells / μl and submitted for single cell long read sequencing.

| Cell line | Provider | Catalog number |
| --- | --- | --- |
| IGROV-1 | NCI DCTD |  |
| SK-OV-3 | ATCC | HTB-77 |
| OVTOKO | JCRB Cell Bank | JCRB1048 |
| OVKATE | JCRB Cell Bank | JCRB1044 |
| OVMANA | JCRB Cell Bank | JCRB1045 |
| COV362 | ECACC | 07071910 Lot# 07G029 |
| COV504 | ECACC | 07071902 Lot# 07I007 |

Reference: ^32^

### Single-cell, long-read library preparation and nanopore sequencing

Approximately 10 ng of cDNA generated from 10x was amplified using the biotinylated version of the forward primer from the ONT protocol, ([Btn]_Fwd_3580_partial_read1_defined) and reverse primer (Rev_PR2_partial_TSO_defined). To get enough cDNA for the pull-down two PCR reactions were carried out using 2X LongAmp Taq (NEB, Cat. M0287S) with the following PCR parameters 94℃ for 3 minutes, with 5 cycles of 94℃ 30 secs, 60℃ 15 secs, and 65℃ for 3 mins, with a final extension of 65℃ for 5 minutes. The cDNA was pooled and cleaned up with 0.8X SPRI ratio with an elution volume of 40μL. Concentration was evaluated using the QuBit HS dsDNA protocol. The amplified cDNA was then captured using 15 μL M270 streptavidin beads (Thermofisher). Beads were washed three times with SSPE buffer (150 mM NaCl, 10 mM NaH_2_PO_4_, and 1 mM EDTA). Beads were then resuspended in 10μL of 5X SSPE buffer (750 mM NaCl, 50 mM NaH_2_PO_4_, and 5 mM EDTA). Approximately 200 ng of the cDNA in 40μL were added together with the 10μL M270 beads and incubated at room temperature for 15 minutes. After incubation, the sample and beads are washed twice with 1mL of 1X SSPE. A final wash is performed with 200 uL of 10 mM Tris-HCl (pH 8.0) and the beads bound to the sample are resuspended 10μL H_2_O. PCR was then performed on-bead using the unbiotinylated version of the primers shown above for 5 cycles according to the same PCR program shown above. A 0.8X SPRI was performed. The cDNA was eluted in 50 μL and concentration was evaluated with QuBit HS dsDNA and Tapestation D5000 DNA kit.

Library preparation for nanopore sequencing was performed according to the LSK-110 kit protocol with the exception of the end-repair step time which was increased to 30 min. 125 fmol of final library was loaded on the PromethION (FLO-PRO002) and sequenced for 72 hr. Reads were basecalled using Guppy v5.0.11.

---

**Figure S1.**
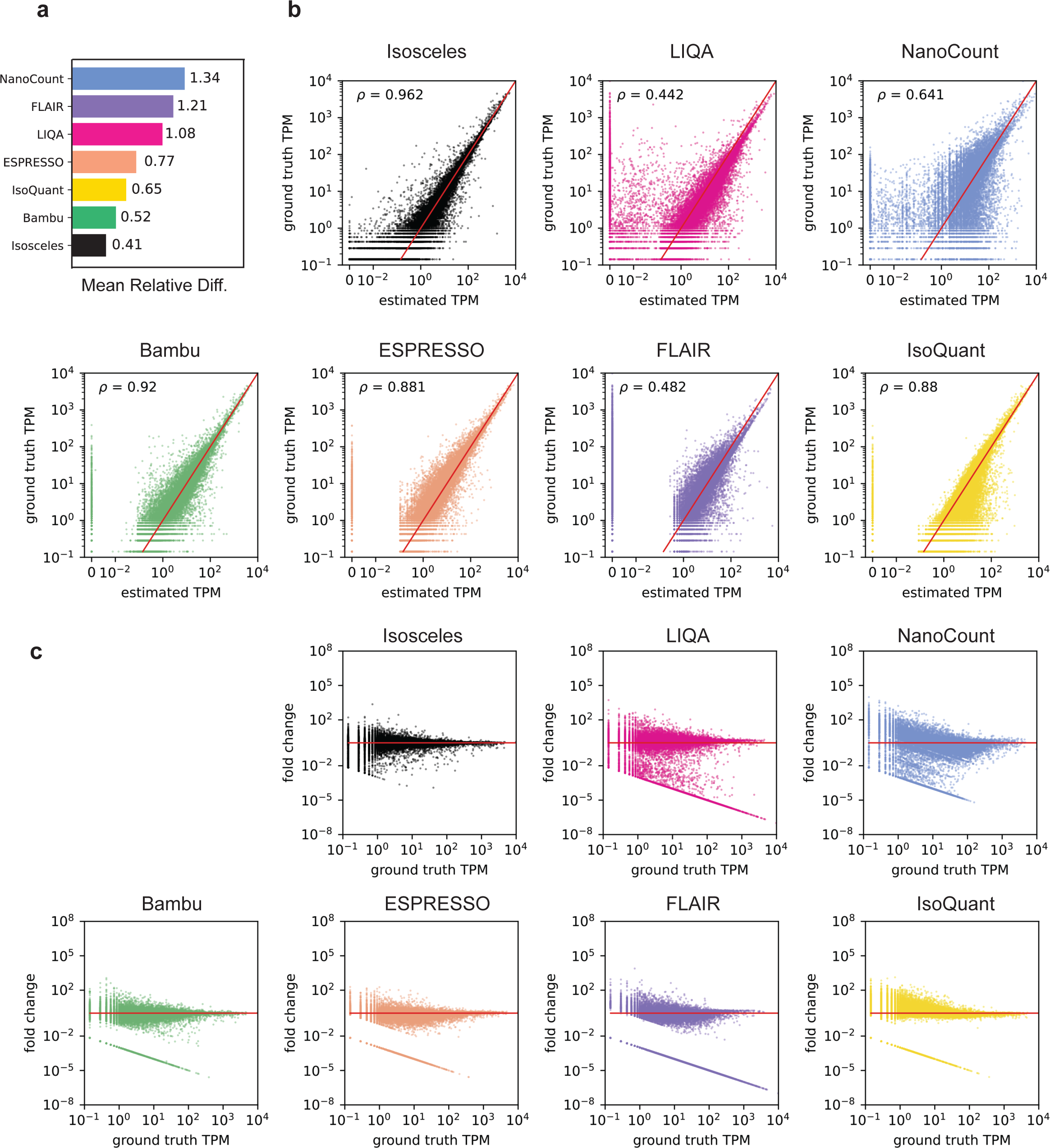
(a) Continued from Fig. 2a, mean relative difference of ground-truth vs. estimated TPM. (b) Scatter plots (with labeled Spearman coefficient) of estimated vs. ground-truth TPM values on log scale. Estimated TPM values below 0.001 are manually assigned a value of 0.001 on the plot. (c) MA plots of the fold change between estimated and ground-truth TPM vs. ground-truth TPM values on log scale. Estimated TPM values below 0.001 are manually assigned a value of 0.001 for the fold change calculation.

**Figure S2.**
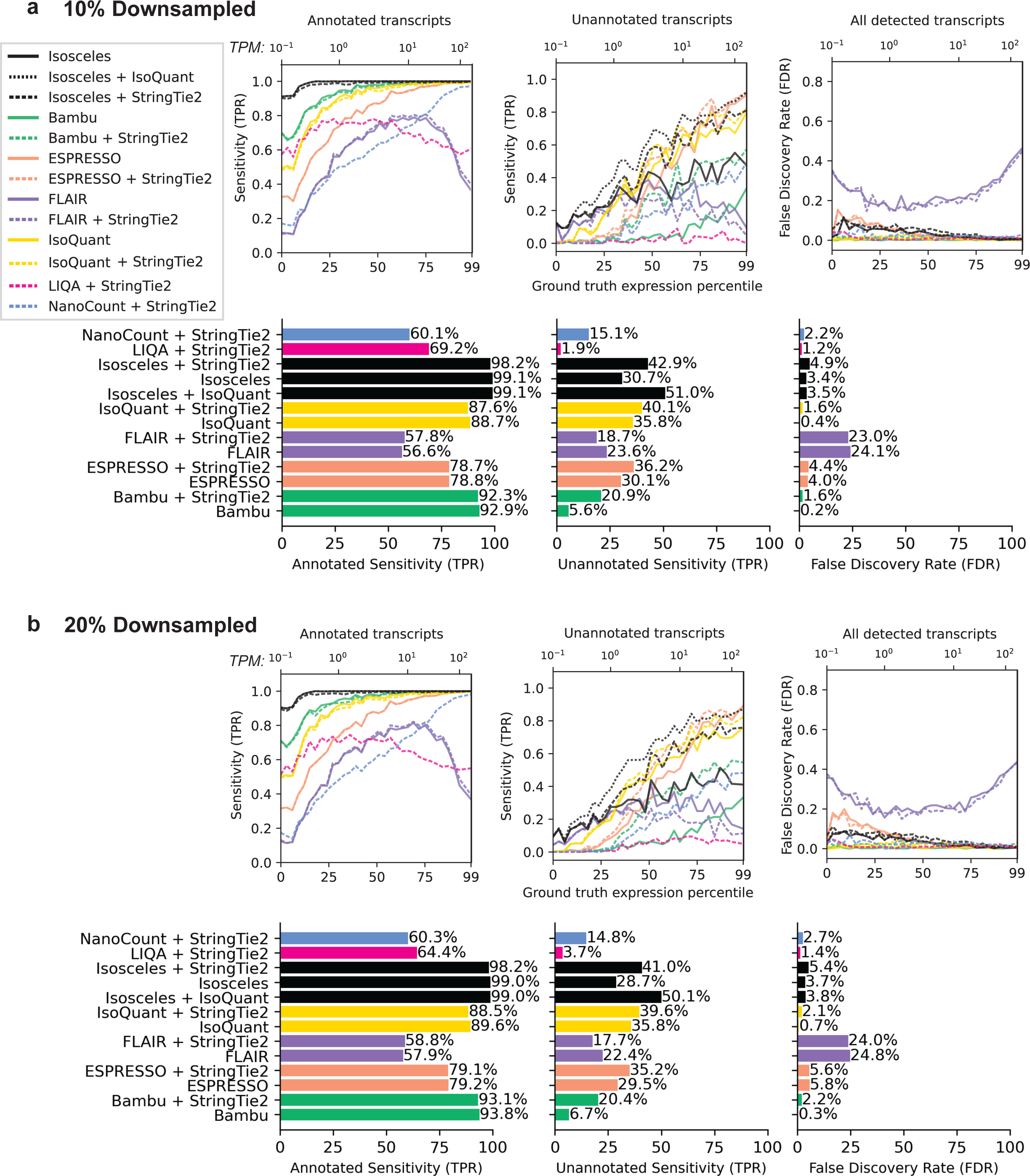
(a, b) Continuation of results from Fig. 2b for 10% and 20% downsampled simulated datasets (see Fig. 2b for additional legend and description).

**Figure S3.**
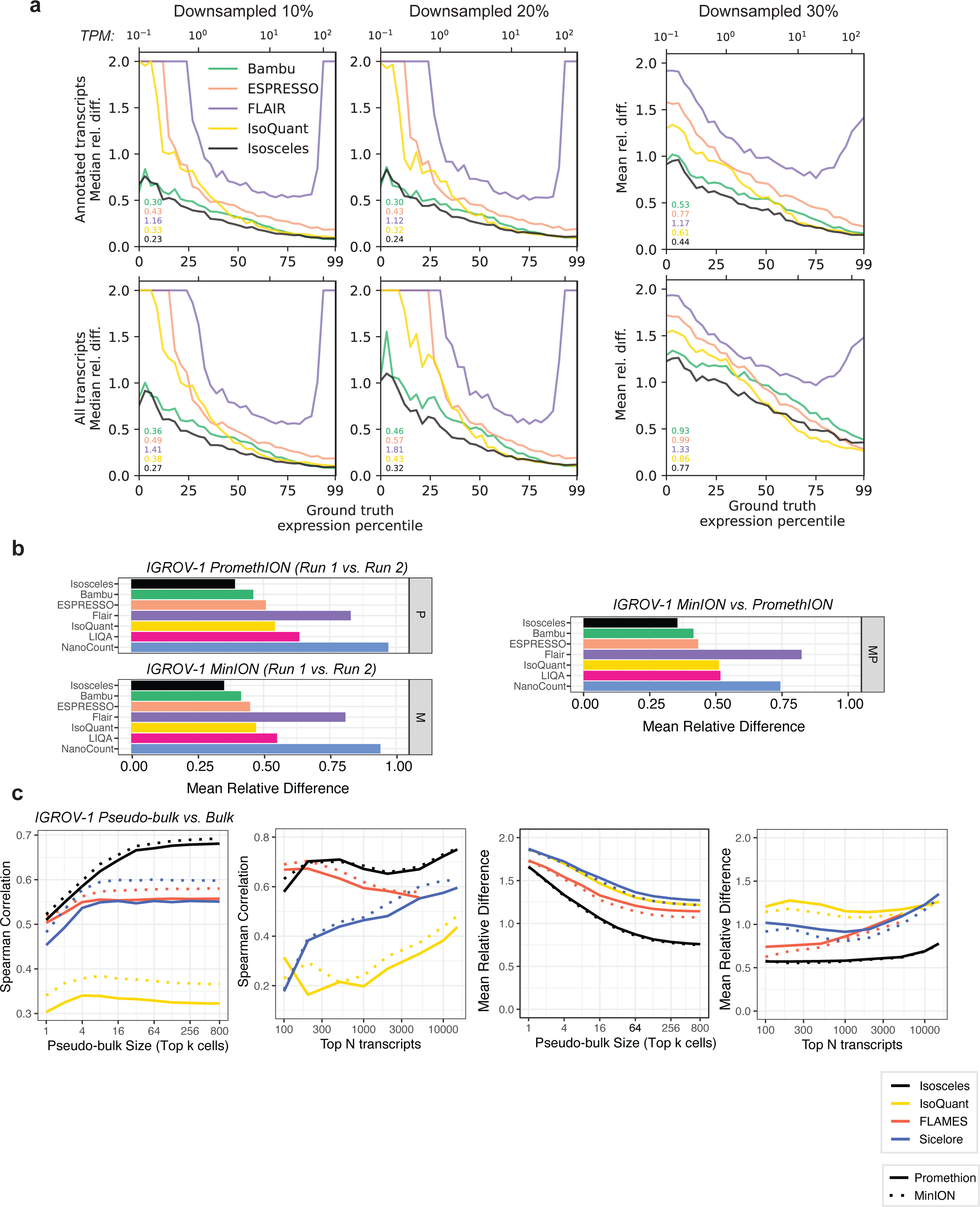
(a) Median relative difference for the additional 10% and 20% downsampling (left; see Fig. 2c for additional legend and description). Mean relative difference for the 30% downsample shown (10% and 20% is concordant, data not shown). (b) Mean relative difference between two runs of the same IGROV-1 sample sequenced with a PromethION (P), MinION (M), or between the two (MP). (c) Correlations and relative difference of pseudo-bulk vs. bulk for top 5000 expressed transcripts in IGROV-1 cells as a function of the number of top ranked cells (by UMI count) included in the pseudo-bulk (left, middle right) and as a function of the top number of transcripts included for the top 64 cells (middle left, right).

**Figure S4.**
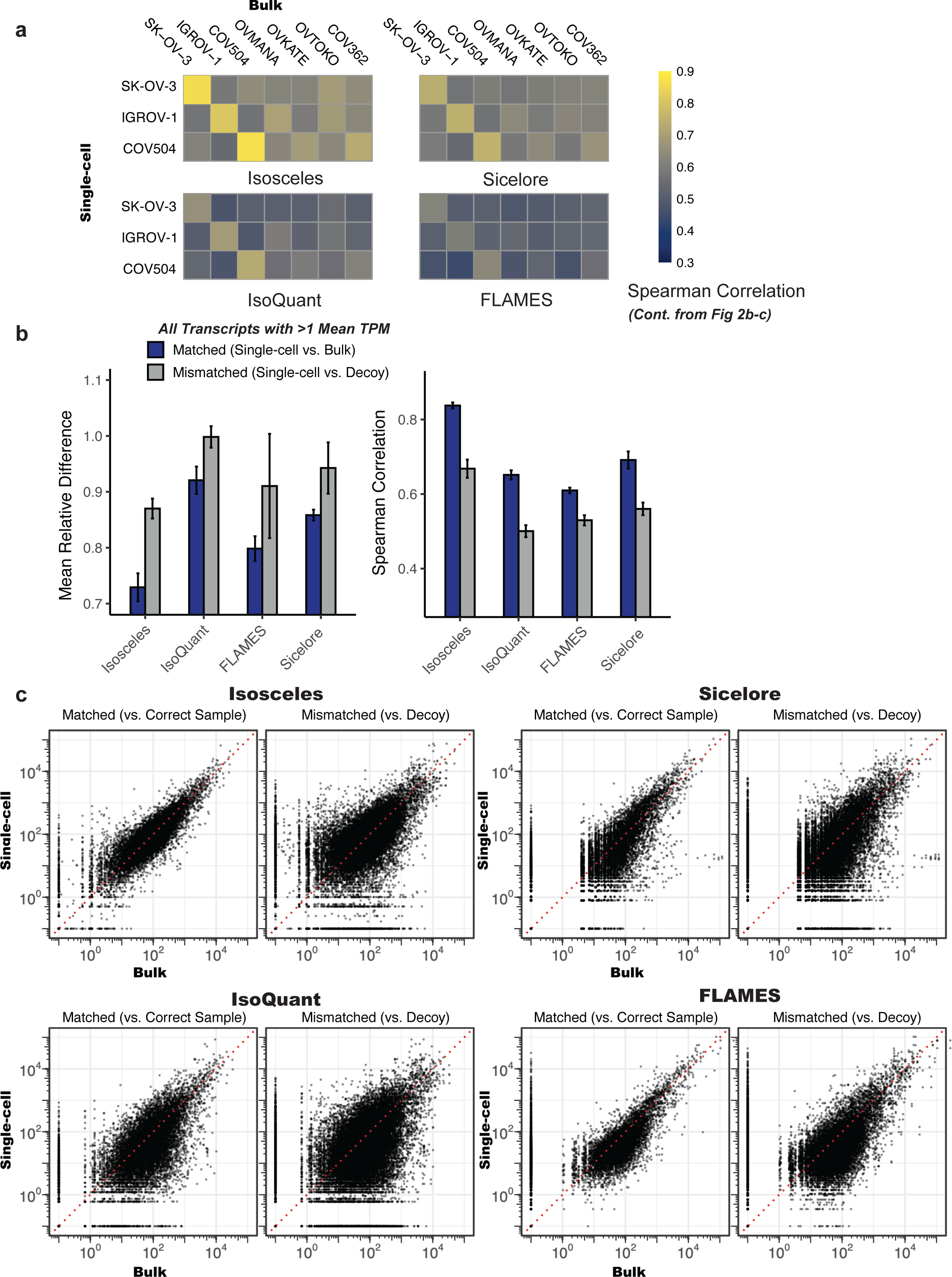
(a) Continued from Fig. 3b, Spearman correlations for each cell line in pseudo-bulk (by genetic identity) vs. the seven bulk nanopore sequenced ovarian cell lines. (b) Alternative version of Fig. 3c, including all transcripts for each program with >=1 TPM among the three cell lines in pseudo-bulk. (c) Continued from Fig. 2c, overlaid scatter plots of all matched (left) and decoy (right) comparisons, where each point is a transcript from one of the comparisons.

**Figure S5.**
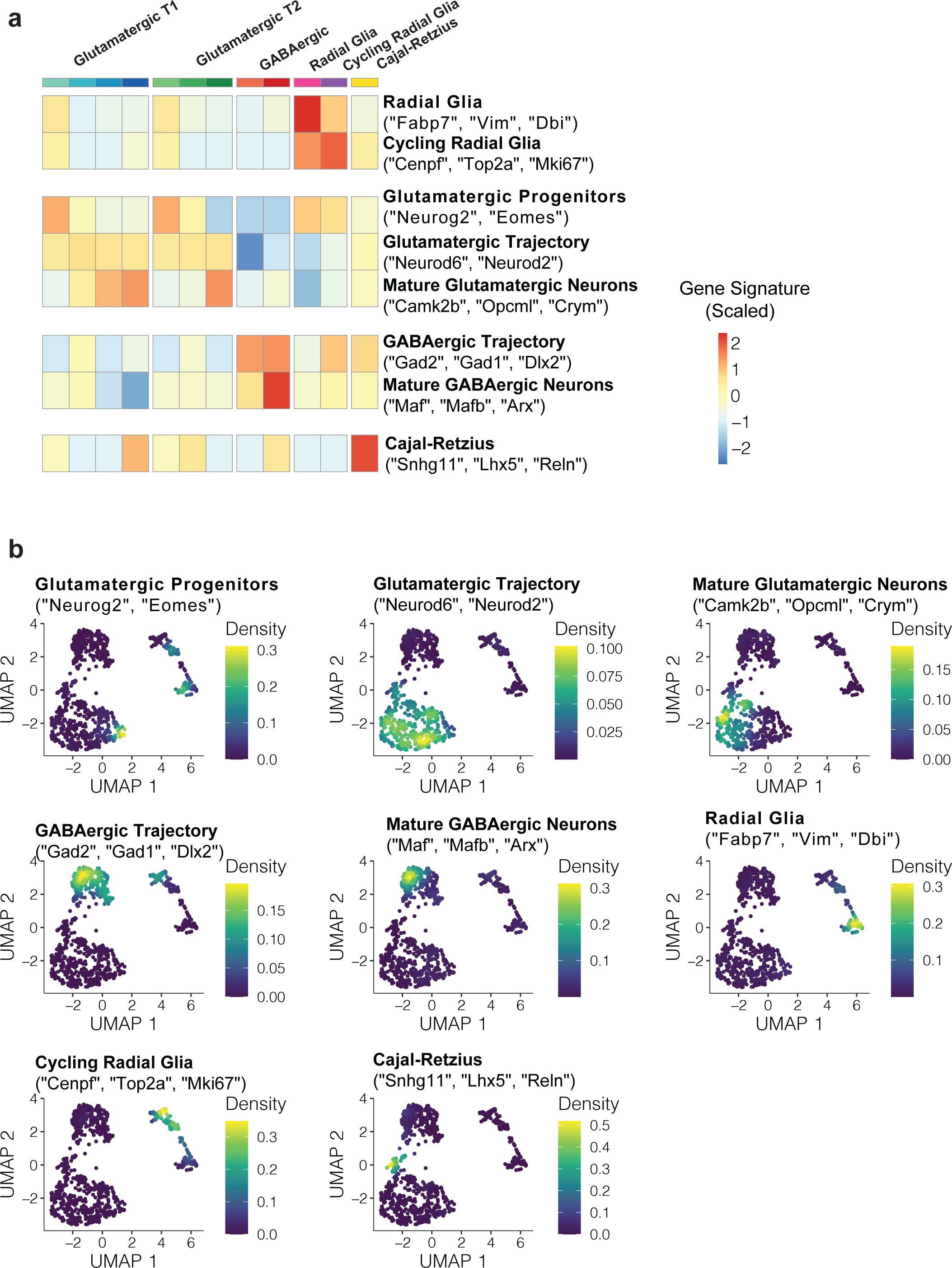
(a) Heatmap of marker gene signature expression along clusters colored according to Fig. 4a. (b) Density plots of each of the corresponding gene signatures overlaid on the UMAP (see Fig 4a).

**Figure S6.**
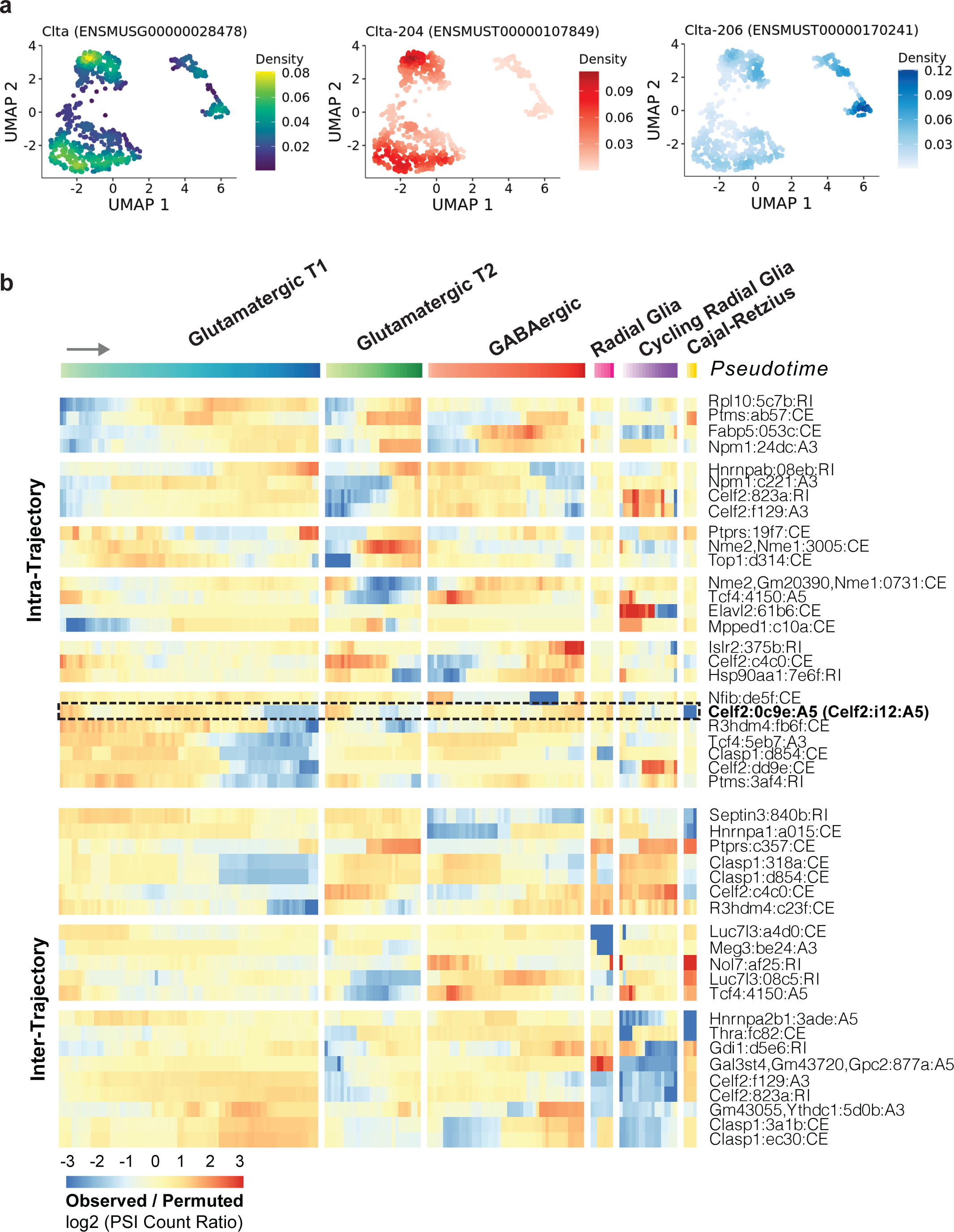
(a) Exemplar result of ‘isoform switching’ analysis, the gene Clta is consistent with the findings highlighted in the original study. The two isoforms 204 and 206 show an expression difference between radial glia and glutamatergic neurons. (b) Expanded version of Fig. 4b plot (with the same scales), but labeled by gene : short hash id : event type. The short hash id matches the ‘psi_event_label’ column label Table S1, which contains the genomic coordinates and Ensembl gene ids. The event type abbreviations are explained in the Methods, eg. CE = Core Exonic interval, A5 = Alternative 5’ splice site, A3 = Alternative 3’ splice site, RI = Retained Intron.

**Figure S7.**
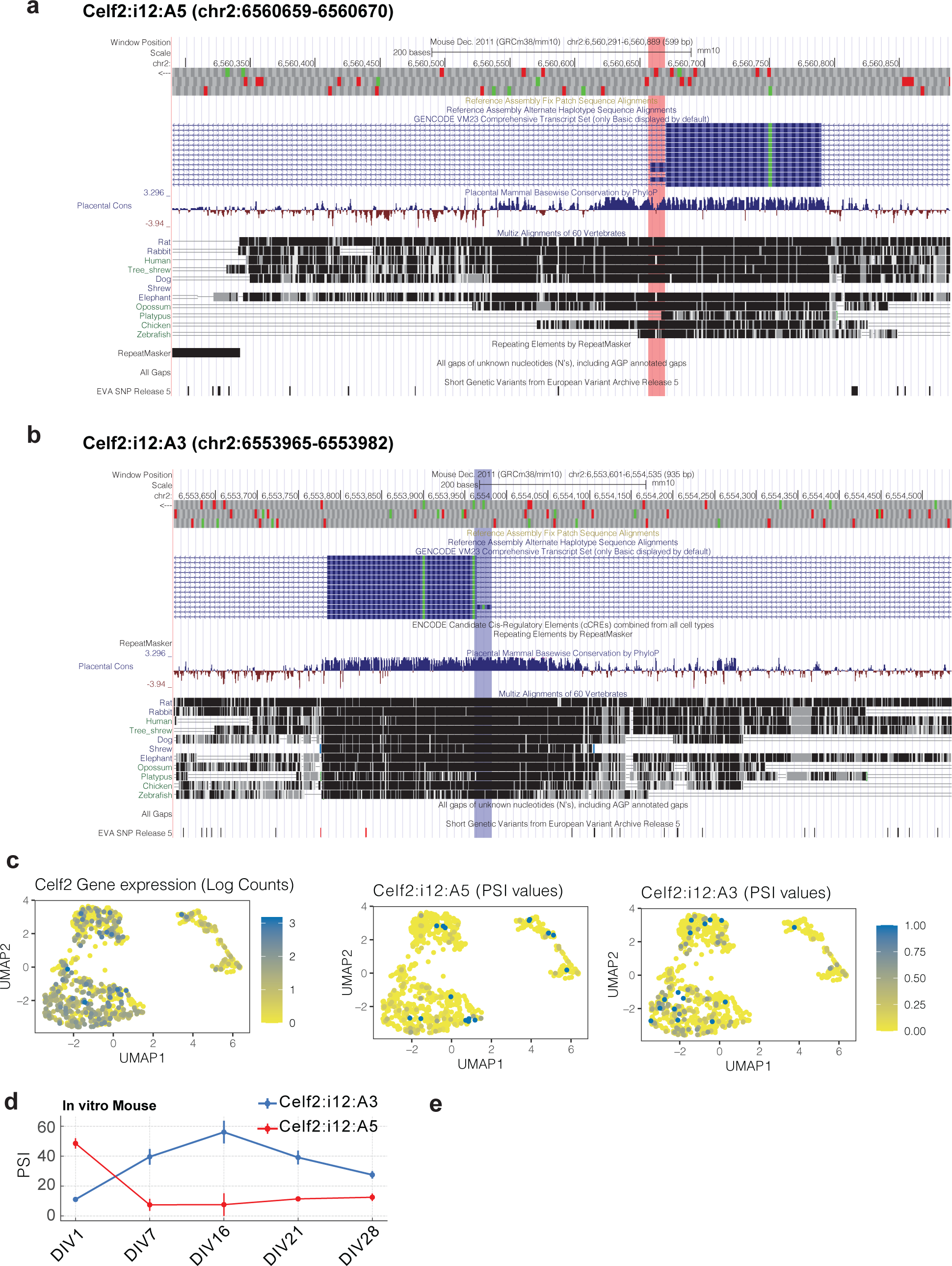
(a) UCSC Genome Browser snapshot of Celf2 intron 12 for the A5 event. Red bar marks the alternative region included by the A5 event, and matches the region marked in red from Fig 4d. (b) Same as panel a but for the A3 event, which matches the blue region in Fig 4d. (c) Extended set of plots matching Fig 4c, but raw gene expression counts, and raw PSI values. (d) PSI values for Celf2:i12:A5 and Celf2:i12:A3 across a mouse in vitro longitudinal glutamatergic neuron differentiation time series ^24,25^.

